# Metabolic Reprogramming by *In Utero* Maternal Benzene Exposure

**DOI:** 10.1101/2020.10.12.336313

**Authors:** Lisa Koshko, Lucas K. Debarba, Mikaela Sacla, Juliana M.B. de Lima, Olesya Didyuk, Patrick Fakhoury, Marianna Sadagurski

## Abstract

Environmental chemicals play a significant role in the development of metabolic disorders, especially when exposure occurs early in life. We have recently demonstrated that benzene exposure, at concentrations relevant to a cigarette smoke, induces a severe metabolic imbalance in a sex-specific manner affecting male but not female mice. However, the roles of benzene in the development of aberrant metabolic outcomes following gestational exposure, remain largely unexplored. In this study, we exposed pregnant C57BL/6JB dams to benzene at 50 ppm or filtered air for 5 days/week (6h/day from gestational day 1 to birth) and studied male and female offspring metabolic phenotypes in their adult life. While no changes in body weight or body composition were observed between groups, 4-month-old male and female offspring exhibited reduced parameters of energy homeostasis (VO_2_, VCO_2_, and heat production). However, only male offspring from benzene-exposed dams were glucose intolerant and insulin resistant at this age. By six months of age, both male and female offspring displayed glucose and insulin intolerance, associated with elevated expression of hepatic gluconeogenesis and inflammatory genes. Additionally, this effect was accompanied by elevated insulin secretion and increased beta-cell mass only in male offspring. Thus, gestational benzene exposure can reprogram offspring for increased susceptibility to the metabolic imbalance in adulthood with differential sensitivity between sexes.

## Introduction

Type 2 Diabetes (T2DM) is an escalating public health concern with complex genetic, behavioral, and environmental origins (Brancati et al. ; Fujimoto ; Shaw et al. 2010). T2DM mostly results from the interaction between genetic, environmental, and other risk factors (Murea et al. 2012). Emerging evidence suggests that environmental chemical exposures may contribute to T2DM by altering whole-body glucose metabolism, insulin resistance, as well as by affecting satiety signals and energy homeostasis in the central nervous system (Kuo et al. 2013). Compelling evidence indicates that exposure to various toxins or air pollution during sensitive periods of early development causes a predisposition to metabolic disorder later in life (Das et al. 2018; Rauh and Margolis 2016; Soh et al. 2018; Suryadhi et al. 2019; Vanker et al. 2018). Specifically, volatile organic compounds (VOCs) have been shown to induce damaging health effects (Dezest et al. 2017; Mohamed et al. 2002; Williams et al. 2019; Yoon et al. 2010).

Benzene is one of the main VOCs found in ambient air pollution (Kliucininkas et al. 2011). Other sources of benzene include processed foods, occupational exposure, vehicle and manufacturing emissions, cigarette smoke, and electronic cigarette fumes (Duarte-Davidson et al. 2001; Minciullo et al. 2014; Pankow et al. 2017; Salviano Dos Santos et al. 2015; Wallace 1989; Williams and Mani 2015). Benzene is known to be carcinogenic at high concentrations, but its association in causing metabolic diseases particularly

T2DM is still emerging. Recent studies demonstrated that benzene non-cancerous health effects at environmental levels include impaired glucose homeostasis and insulin resistance (Abplanalp et al. 2017; Amin et al. 2018; Debarba et al. 2020). We have recently shown that chronic exposure of adult mice to benzene at the levels relevant to human exposures associated with cigarette smoke (50 ppm) induces severe metabolic imbalance associated with central hypothalamic inflammation and endoplasmic reticulum (ER) stress in a sex-specific manner (Debarba et al. 2020). This data indicates that exposure to benzene can affect an individual’s susceptibility to metabolic imbalance and the development of T2DM.

Most epidemiological studies investigating the association between benzene exposure and insulin resistance have focused on young adult or elderly populations living in highly polluted areas (Abplanalp et al. 2017; Burg and Gist 1998; Choi et al. 2014). However, very few studies have assessed the risk of benzene exposure during pregnancy on the long-term metabolic health of the grown offspring. Recent epidemiological studies have highlighted the association of benzene exposure with increased risk for gestational diabetes (Williams et al. 2019). Similarly, epidemiological data have shown a positive association between cigarette smoke exposure *in utero* and increased risk of obesity and metabolic imbalance in offspring (Behl et al. 2013). A study of traffic pollutants (PM10 and benzene) in a community in northern Italy showed an association between exposure to PM10 and different birth defects (De Donno et al. 2018). However, the direct link between maternal exposure to benzene during pregnancy and metabolic disease in offspring is not clear.

The current study sought to determine whether exposure of pregnant females to benzene alters the susceptibility of the young and/or adult offspring to metabolic imbalance. Pregnant females were exposed to filtered air or concentrations of benzene at the levels relevant to cigarette smoke throughout the gestation by direct inhalation for 6 hours per day, 5 days a week. Under these conditions, benzene exposure had no significant effect on litter size or composition and no significant changes in animals white blood cell counts (Debarba et al. 2020). Our findings indicate that maternal exposure to benzene induces deleterious effects on energy expenditure, glucose metabolism, and insulin resistance in adult offspring in a sex-specific manner, providing the evidence that gestational benzene exposure is a potential risk factor for metabolic disorders.

## Methods

### Animals

8-9 weeks-old male and female C57BL/6 were purchased from The Jackson Laboratory. After a one-week habituation period, female and male mice were mated, and mice were considered pregnant if a vaginal plug was observed. In the present study, we used 5-6 pregnant dams on gestation day 0 (GD0). During the gestational period of 21 days, the pregnant dams were exposed 6 hours per day, 5 days a week to benzene [50 ppm] in the inhalation chamber (see below). Pups were delivered outside of the chamber. Females were fed *ad libitum* throughout pregnancy and lactation. At the age of 21 days, pups were weaned onto a chow diet (Purina Lab Diet 5001). Most of the presented data related to both male and female mice unless otherwise stated.

### Benzene exposure

Pregnant dams in inhalation chambers using FlexStream™ automated Perm Tube System (KIN-TEK Analytical, Inc) were exposed to benzene concentration of 50 ppm for 6h/day, 5 days a week as before (Debarba et al. 2020). FlexStream™ automated Perm Tube System allows creating precision gas mixtures. This unit provides a temperature-controlled permeation tube oven, dilution flow controls, and front panel touch-screen interface. Mixtures are produced by diluting the miniscule flow emitted from Trace Source™ permeation tubes with a much larger flow of inert matrix gas, typically nitrogen or zero air. Control dams were breathing regular filtered room air. All mice were provided with water ad libitum and housed in temperature-controlled rooms on a 12-hour/12-hour light-dark cycle.

### Metabolic Analysis

Lean and fat body mass were assessed by a Bruker Minispec LF 90II NMR-based device. Intraperitoneal glucose tolerance test (GTT) was performed on mice fasted for 5-6 hours. Animals were then injected intraperitoneally with D-glucose (2 g/kg) and blood glucose levels were measured as before (Sadagurski et al. 2010). For an insulin tolerance test (ITT), animals fasted for 4 hours received an intraperitoneal injection of insulin (Humulin R, 0.8 U/kg) diluted in sterile saline. Blood glucose concentrations were measured at the indicated time points. Blood insulin levels were determined on serum from tail vein bleed using a Mouse Insulin ELISA kit (Crystal Chem. Inc.). In all tests, tail blood glucose levels were measured at the indicated times after injection. Metabolic measurements of energy homeostasis were obtained using an indirect calorimetry system (PhenoMaster, TSE system, Bad Homburg, Germany). The mice were acclimatized to the cages for 2 days and monitored for 5 days while food and water were provided *ad libitum.*

### RNA extraction and qPCR

Total RNA was extracted from the liver with Trizol (Gibco BRL) and 1000 ng of RNA was used for cDNA synthesis using iscript cDNA kit (Bio-Rad Laboratories Inc.). Quantitative real-time PCR was performed using the Applied Biosystems 7500 Real-Time PCR System. The following SYBER Green Gene Expression Assays (Applied Biosystems) were used in this study. See Table 1 for primers sequence. Each PCR reaction was performed in triplicate. Lack of reverse transcription (-RT) was used as a negative control for contaminating DNA detection. Water instead of cDNA was used as a negative control, and the housekeeping gene β-actin was measured in each cDNA sample. Gene transcripts in each sample were determined by the ΔΔCT method. For each sample, the threshold cycle (CT) was measured and normalized to the average of the housekeeping gene (ΔCT). The fold change of mRNA in the unknown sample relative to the control group was determined by 2-ΔΔCT. Data are shown as mRNA expression levels relative to the control group.

### Statistical analysis

Data sets were analyzed for statistical significance using Statistica software (version 10). An analysis of variance (ANOVA) with repeated measurements was used to analyze body weight, GTT and ITT tests. Additionally, an analysis for a two-tail unpaired Student’s t-test was used when two groups were compared. The level of significance (α) was set at 5%.

## Results

### Gestational benzene exposure does not alter body weight in the offspring

We exposed pregnant females to benzene at the concentration of 50 ppm throughout the pregnancy using an exposure chamber for 6 hr/day, 5 days/wk for 21 days (Debarba et al. 2020). Figure 1A illustrates a schematic timeline of the experimental design. This adapted regimen for benzene inhalation resembles chronic human benzene exposure levels in highly polluted gas stations, or exposure to tobacco smoke (De Donno et al. 2018; Lorkiewicz et al. 2019; Pankow et al. 2017). We have recently demonstrated that under these conditions, benzene exposure has no significant changes in white blood cell counts in benzene exposed animals versus control animals breathing room air (Debarba et al. 2020). Maternal exposure to benzene did not significantly affect male and female offspring body mass at weaning and throughout adulthood (Figure 1B and C). Additionally, the lean and fat body mass of male and female offspring were not affected by maternal benzene exposure (Figure 1D and E).

**Figure 1.**
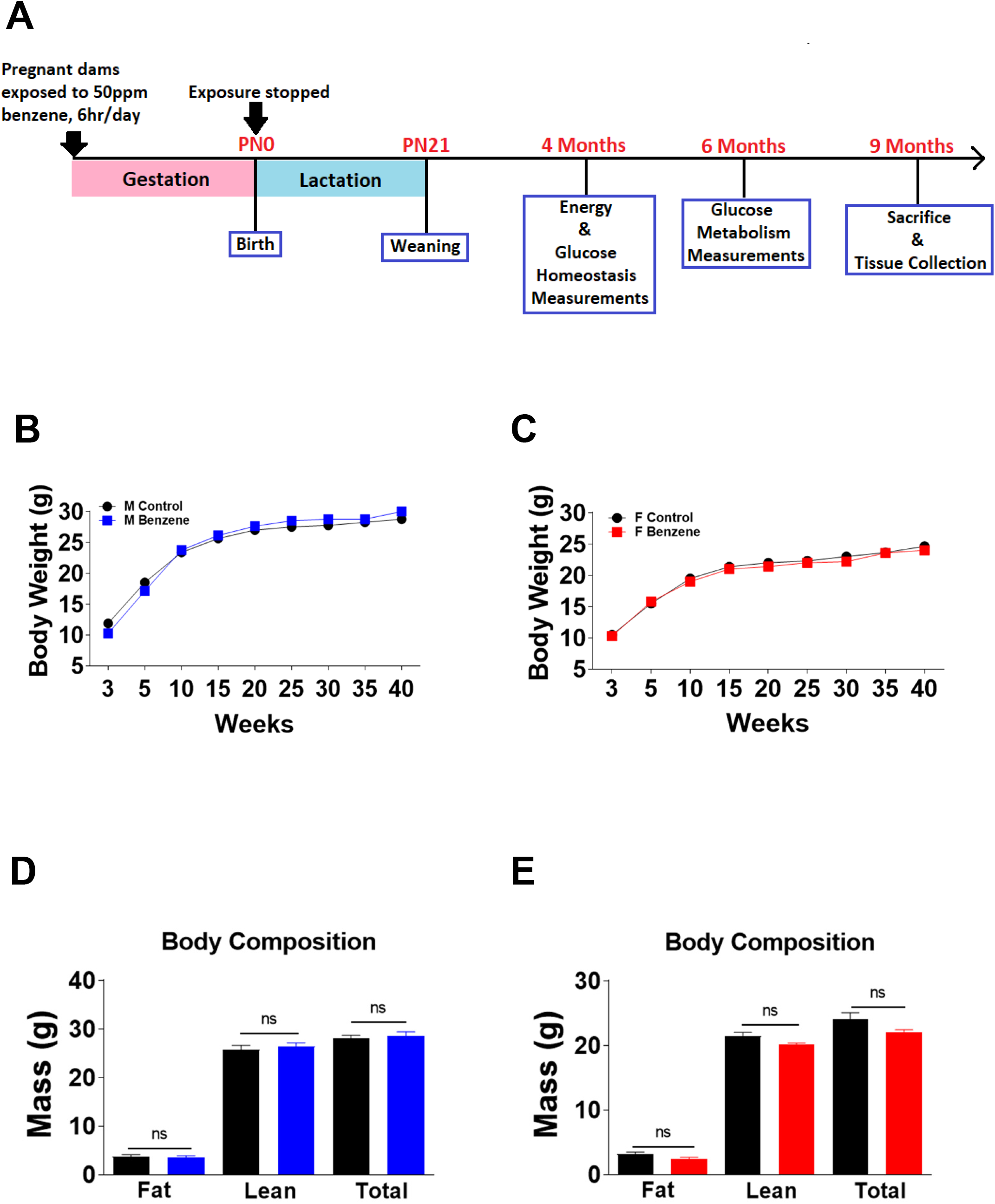
Gestational benzene exposure does not alter body weight in the offspring. (**A**) Experimental design timeline. Effect of gestational benzene exposure on body weight gain (g) of males (**B**) and females (**C**); fat mass (g), lean mass (g) and total mass (g) of males (**D**) and females (**E**) measured by magnetic resonance imaging (echo MRI), data are expressed as the mean ± SEM (n= 6). Repeated measures ANOVA were further analyzed with Newman-Keuls post hoc analysis or t-test if necessary to compare between only two groups of the predictor variable.

### Gestational benzene exposure predisposes to impaired energy expenditure and glucose metabolism in offspring

We next assessed whole-body energy homeostasis in young adult male and female offspring (Figure 2-3). Both male and female offspring of benzene-exposed dams demonstrated significantly decreased oxygen consumption (VO_2_), carbon dioxide (VCO_2_) production and heat production during the 24h period (p < 0.01, Figure 2A-C and 3A-C). Interestingly, male offspring of benzene-exposed dams exhibited a greater reduction in energy expenditure parameters compared to female offspring as indicated by stronger reduction in VO_2_, VCO_2_ and heat production during both light and dark cycles (p < 0.01, Figure 2A-C and 3A-C). There was no significant difference in locomotor activity levels or food intake among the groups (Figure 2D, 3D, and data not shown).

**Figure 2.**
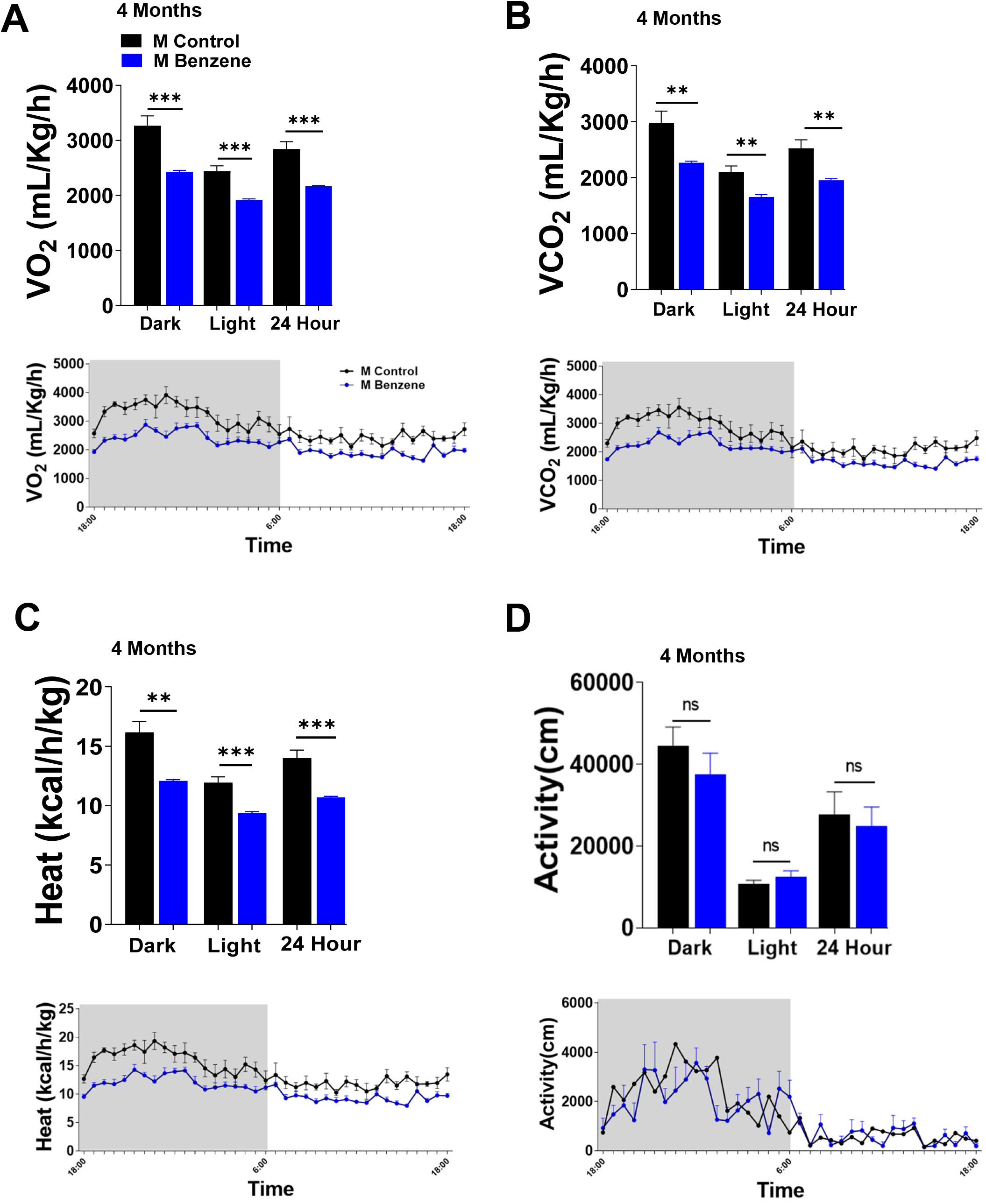
Gestational benzene exposure impairs energy homeostasis in 4-month-old male offspring. Effect of gestational benzene on (**A**) oxygen consumption (VO_2_ – mL/kg/h); (**B**) carbon dioxide production (VCO_2_-mL/kg/h); (**C**) heat production (kcal/h/kg); (**D**) Locomotor activity during light, dark, and the entire 24-h cycles of male offspring at 4-months of age. The line graphs depict variations throughout each cycle. Data are expressed as the mean ± SEM (n= 5). Repeated measures ANOVA were further analyzed with Newman-Keuls post hoc analysis or t-test if necessary, to compare between only two groups of the predictor variable, (* =*vs* control; \**p* < 0.05; \*\**p* <0.01).

**Figure 3.**
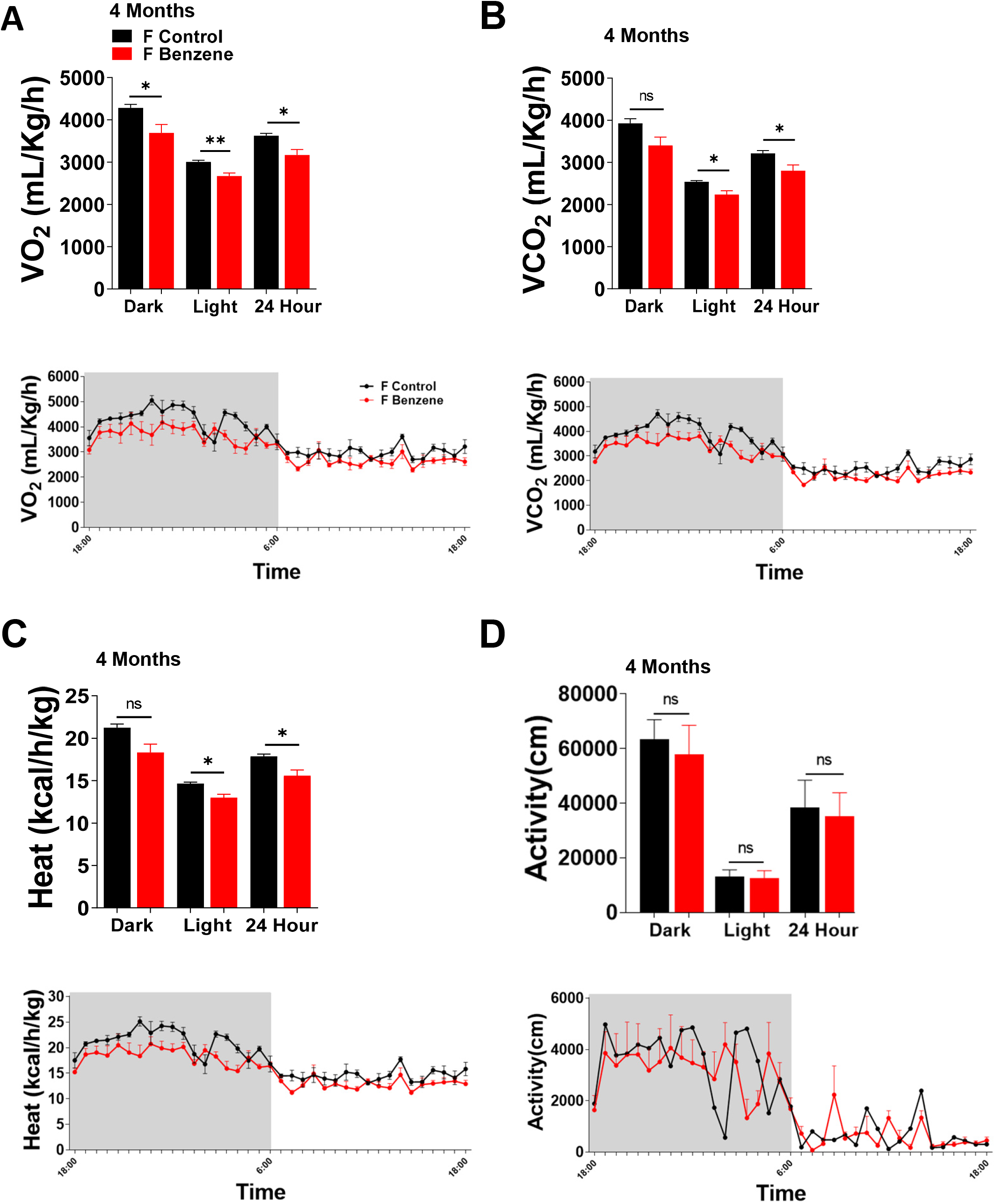
Gestational benzene exposure impairs energy homeostasis in 4-month-old female offspring. Effect of gestational benzene on (**A**) oxygen consumption (VO_2_ – mL/kg/h); (**B**) carbon dioxide production (VCO_2_-mL/kg/h); (**C**) heat production (kcal/h/kg); (**D**) Locomotor activity during light, dark, and the entire 24-h cycles of male offspring at 4-months of age. The line graphs depict variations throughout each cycle. Data are expressed as the mean ± SEM (n= 5). Repeated measures ANOVA were further analyzed withNewman-Keuls post hoc analysis or t-test if necessary, to compare between only two groups of the predictor variable, (* =*vs* control; \**p* < 0.05; \*\**p* <0.01).

Exposure of male mice to benzene during gestation resulted in severe hyperglycemia, as indicated by glucose tolerance test (Figure 4A), and by significantly higher area under the curve (AUC) and HOMA-IR, an index of insulin resistance (Figure 4 A, C, and D) in 4-month-old male offspring. On the other hand exposure of female mice to benzene during gestation had no effect on glycemia at this age (Figure 4B). However, by six-month of age both male and female offspring of benzene-exposed dams exhibited marked glucose intolerance (Figure 5A and B). By this age male offspring of benzene-exposed dams developed severe insulin resistance (p<0.05), while female offspring, demonstrated a slight impairment in insulin tolerance (p<0.05, Figure 5C, and 5D). To examine glucose-stimulated insulin secretion in six-month-old male offspring, we measured plasma insulin at baseline, 15 min, and 30 min after i.p. glucose challenge (Figure 6A). While insulin fasting levels were not different between the groups, the overall glucose-stimulated insulin secretory response was significantly increased in the male offspring of benzene-exposed dams (p<0.05 vs control offspring). Additionally, maternal benzene exposure modulated islet morphology in these male offspring and they were characterized by a significant increase in β-cells mass compared to control male offspring (Figure 6B and 6C). Notably, β-cell mass in female offspring of benzene exposed dams was similar to control female offspring (Supplementary Figure 1), confirming the observed sex differences in metabolic adaptation to benzene exposure (Debarba et al. 2020).

**Figure 4.**
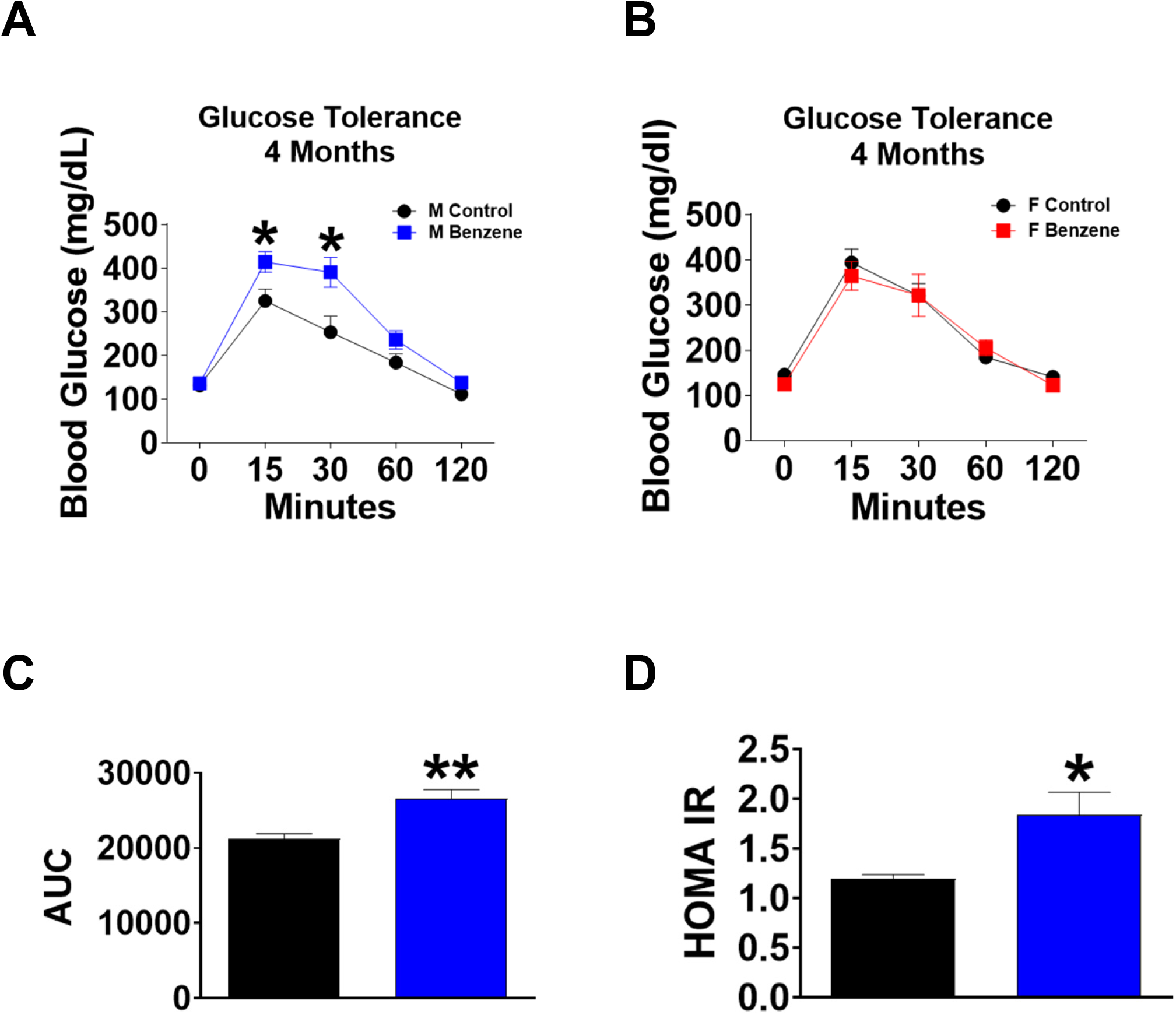
Gestational benzene exposure impairs glucose metabolism in the offspring in a sex-specific manner at 4 months of age. Effect of gestational benzene on glucose tolerance test (GTT) of (**A**) male and (**B**) female offspring; (**C**) Area under the curve (AUC); (**D**) HOMA – IR of male offspring, at 4 months of age. Data are expressed as the mean ± SEM (n=6 - 7). Repeated measures ANOVA were further analyzed with Newman-Keuls post hoc analysis or by t-test if necessary to compare between only two groups of the predictor variable, (* p < 0.05 vs control).

**Figure 5.**
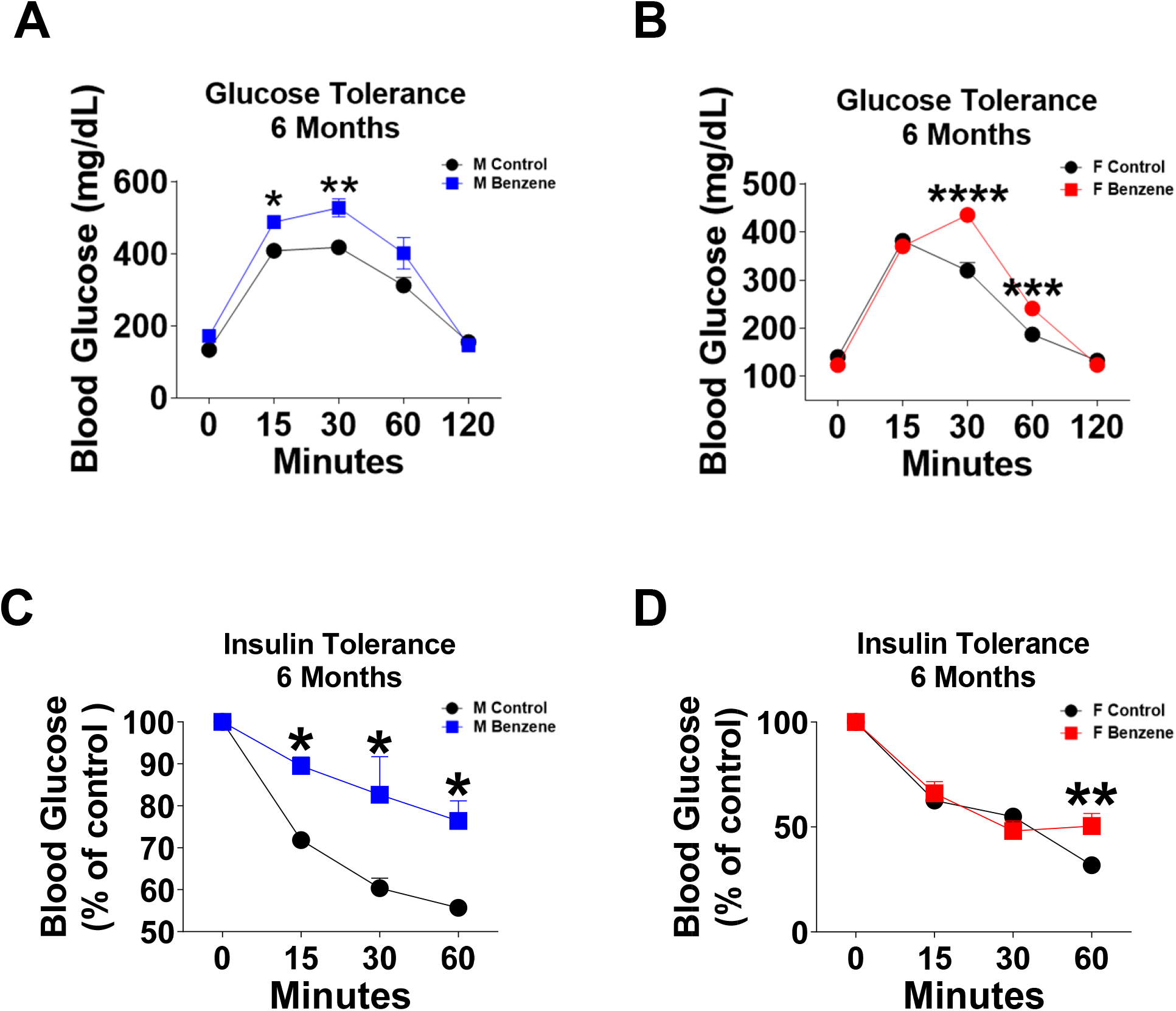
Gestational benzene exposure impairs glucose and insulin tolerance in the offspring at 6 months of age. Effect of gestational benzene on GTT of (**A**) male and (**B**) female offspring; Insulin tolerance test (ITT) of (**C**) male and (**D**) female offspring. Data are expressed as the mean ± SEM (n= 6 - 7). Repeated measures ANOVA were further analyzed with Newman-Keuls post hoc analysis, (* p < 0.05 vs control).

**Figure 6.**
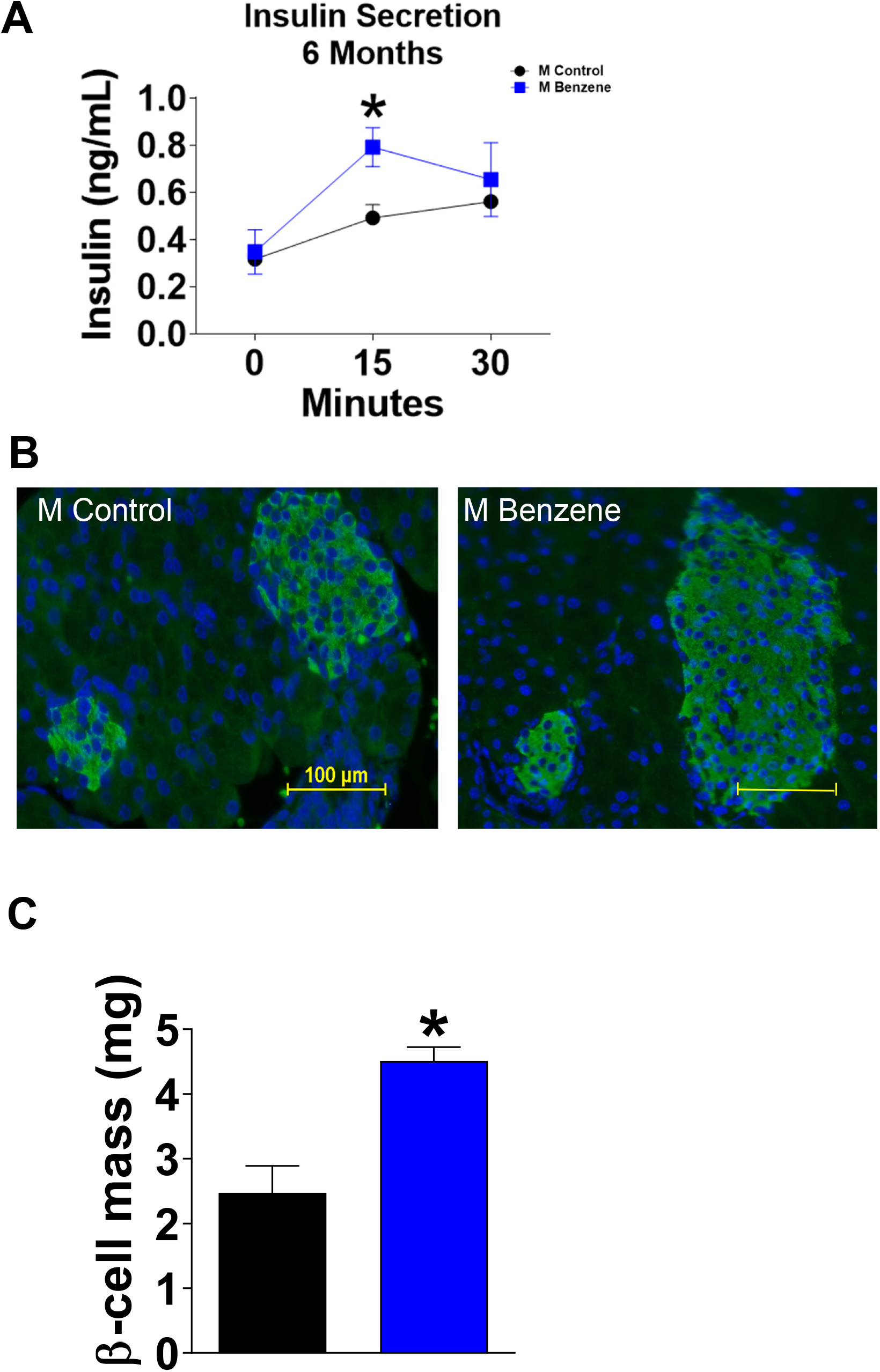
Gestational benzene exposure impairs insulin secretion and beta cells mass in male offspring. (**A**) Insulin secretion; (**B**) Immunofluorescence staining for insulin (green) and DAPI (blue) in representative pancreatic sections from 9-month-old male offspring. **(C)** Quantitation of β-cell mass in male offspring. Data are expressed as the mean ± SEM (n= 6 - 7). (* p < 0.05 vs control).

To further investigate the potential mechanisms of metabolic imbalance in benzene exposed offspring, we analyzed the mRNA levels of genes associated with gluconeogenesis, lipid metabolism, and inflammation in the livers from male and female offspring (Figure 7). Strikingly, we observed a significant increase in hepatic inflammatory genes *Ikk, TNFa,* and *Il6* in both nine-month-old male and female offspring of benzene-exposed dams, indicating that gestational benzene exposure induced hepatic inflammation in the offspring (Figure 7A and 7C). Additionally, both male and female offspring exhibited an increase of - C/EBP homologous protein (*Chop*), a gene involved with the ER-stress pathway (Figure 7A and 7C). We also detected an increase in the expression of genes associated with gluconeogenesis (*Pepck and Gck*) in livers from benzene exposed offspring as compared to control offspring mice (Figure 7B and 7D). No differences were observed in some additional genes associated with inflammation and ER stress, such as *IL1, Emr1, Ire1a*, and *Xbp1s*, or genes associated with lipid and fatty acids synthesis (Supplementary Figure 2). Altogether, these results reveal that benzene exposure during gestation can greatly impair glucose homeostasis and energy balance in adult offspring predisposing them to the development of the metabolic syndrome.

**Figure 7.**
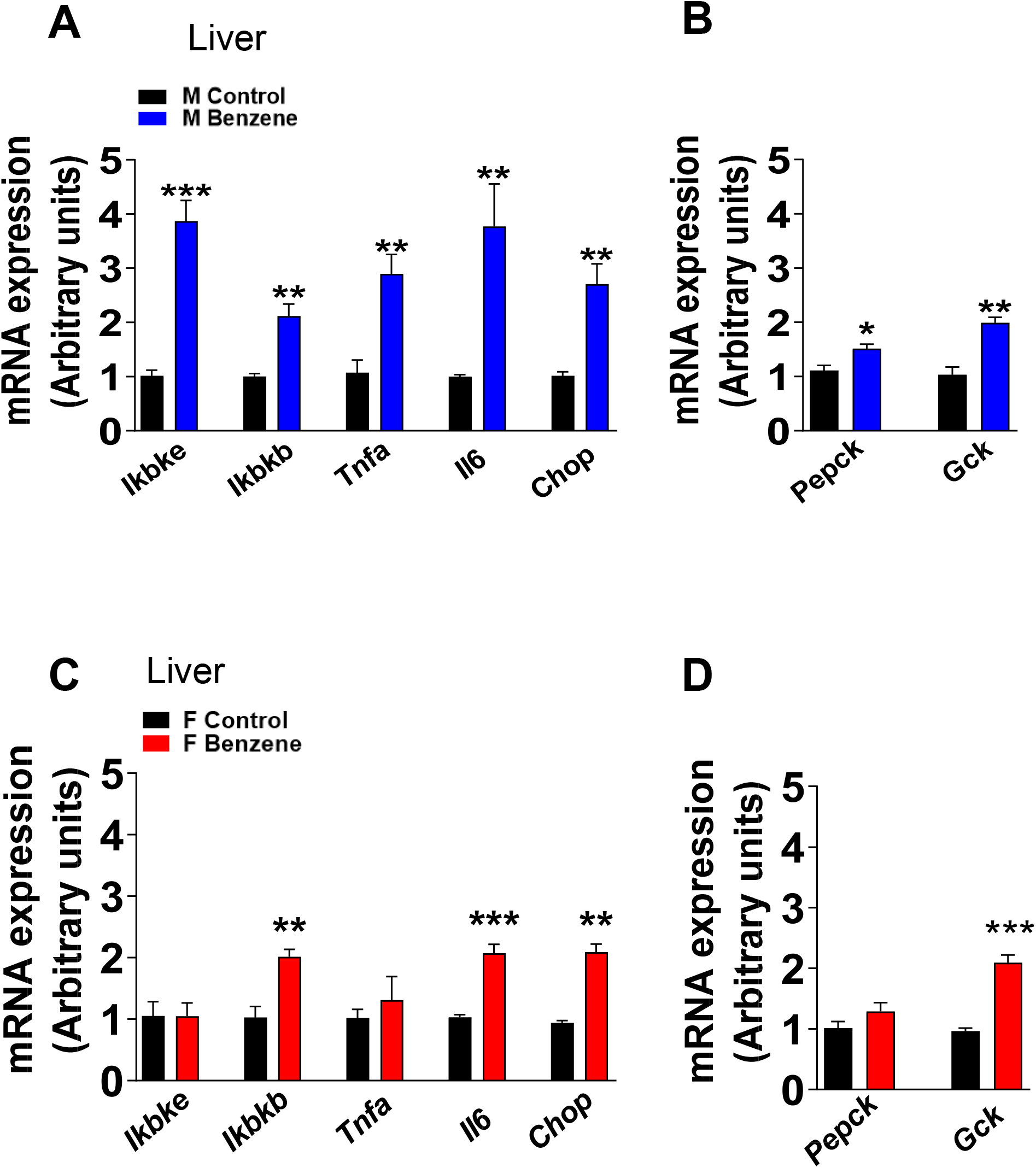
Gestational benzene exposure alters the hepatic gene expression of offspring. qPCR of hepatic inflammatory and ER stress genes (*Ikbkb, Ikbke, TNFa, Il6, CHOP*) of (**A**) males and (**C**) females. qPCR of hepatic gluconeogenesis genes (*PEPCK, Gck*) of (**B**) males and (**D**) females. Data were expressed as the mean ± SEM (n = 4 −5) and analyzed by t-test (* =*vs* control; \**p* < 0.05, **p < 0.01; ***p < 0.001).

## Discussion

This study is the first to demonstrate the severe whole-body metabolic imbalance in young adult offspring born to dams exposed to benzene during pregnancy at concentrations relevant to smoking. Metabolic imbalance in offspring was independent of body weight changes and was associated with impaired energy homeostasis and glucose metabolism, affecting both sexes, although the effect was more severe in adult male offspring. Our data provide a link between early-life exposure to an environmental pollutant and the risk for developing metabolic syndrome later in life (in offspring). Further, it allows elucidating some of the risk factors for metabolic syndrome that pose burden at the population level.

Benzene has been listed as an endocrine-disrupting chemical, but its association with the development of childhood metabolic disorders has not been studied in detail, especially when exposure occurs early in life. We show that gestational benzene exposure has no effect on the offspring’s body weight or body mass at weaning and up to 40 weeks, however, it is an indicator of insulin resistance and metabolic imbalance in adulthood. Adult male offspring exposed to benzene during gestation developed hyperglycemia and severe insulin resistance followed by increased β-cell mass and elevated expression of hepatic gluconeogenic genes. The effect of gestational benzene exposure on female offspring metabolic became apparent only by six-months of age, presented by hyperglycemia and mild insulin resistance with normal β-cell function. Both male and female offspring exhibited a considerable increase in hepatic gene expression associated with inflammation and ER stress. Interestingly, we have recently demonstrated that adult female mice are completely resistant to the negative metabolic consequences of chronic benzene exposure that affects only male mice (Debarba et al. 2020). Benzene crosses the placenta and can be found in the placenta and fetuses immediately following exposure, exerting its effects on developing tissues and cells (Dowty et al. 1976). As such, our data identify gestation as a critical period for adult metabolic perturbations, with increased sensitivity of male offspring to the toxic effects of gestational benzene exposure. In contrast, the protective sex-specific effects in adult females observed in our recent study may have a hormonal basis (Piazza and Urbanetz 2019); however, a causal relationship for these observations remains to be elucidated.

Glucose intolerance and insulin resistance displayed in offspring from benzene-exposed dams indicate that maternal benzene exposure has a strong influence on the offspring’s glucose metabolism likely by predisposing fetal organs involved in glucose disposal and insulin sensitivity to metabolic imbalance. Insulin secretion following glucose load was significantly higher in insulin-resistant male offspring from benzene exposed dams, suggesting an adaptive β-cell response to compensate for insulin resistance, which is in line with increased β-cell mass in these animals. Interestingly, cigarette smoking during gestation induced changes in β-cell function including increased β-cell apoptosis and decreased β-cell mass in rodents (Bruin et al. 2007; Holloway et al. 2005). Islet changes and poor pancreaticβ-cell viability were previously reported due to benzene exposure in rodents (Bahadar et al. 2015). The pancreas is considered to be especially susceptible to oxidative stress-mediated tissue damage (Lenzen et al. 1996). β-cell hypertrophy and hyperplasia occur during β-cell compensation to increase β-cell mass in response to hyperglycemia in diabetogenic states (Cerf et al. 2012). Similarly to insulin resistant and diabetic mice models, increased β-cell mass in male offspring can be likely attributed to β-cell hypertrophy and hyperplasia (Jones et al. 2010).

Both male and female offspring from benzene exposed dams exhibited reduced parameters of energy homeostasis, including O_2_, CO_2_, and heat production. These findings suggest that gestational benzene exposure could alter the hypothalamic regulation of energy homeostasis in both sexes (Bouret and Simerly 2004). While nine-month-old male and female offspring display normal body weight, despite reduced energy and heat production, it is reasonable to hypothesize that during the aging process, such metabolic imbalance will ultimately affect the feeding and weight of these animals. In rodents, the hypothalamic energy balance-regulating system is structurally and functionally immature at the start of postnatal life, and the plasticity of the hypothalamic circuitry provides a route by which environmental signals can regulate energy homeostasis (Bouret 2009; 2010). Our recent findings indicate that benzene exposure induces severe hypothalamic inflammation and ER stress in a sex-specific manner (Debarba et al. 2020). Perinatal exposure to persistent organic pollutants (POPs) reduces energy expenditure with an associated increase in body weight in adult female offspring (La Merrill et al. 2014). In this regard, various studies have demonstrated the sex-specific effect of perinatal exposure to different pollutants and changes in energy homeostasis in the offspring (Heindel et al. 2017). Further research is needed to address the role of hypothalamic “programming” in response to gestational benzene exposure in both sexes.

Taken together, these data indicate that benzene exposure during gestation is the critical window to induce metabolic syndrome in the adult offspring, where male offspring are more sensitive to metabolic imbalance. Gestational exposure to benzene can interfere with epigenetic programming of gene regulation, thereby influencing the risk of diabetes development later in life. Thorough studies of genome-wide analysis will be needed to understand the epigenetic changes triggered by gestational benzene exposure. Such studies can improve our understanding of the possible maternal factors, underlying the programming of metabolic adversity across the life course of the offspring and identify the critical developmental period during which maternal exposure programs altered metabolic control in the offspring.

## Acknowledgments

The authors thank Madelynn Koch and Li Mao for technical assistance.

---

**Supplementary Figure 1.**
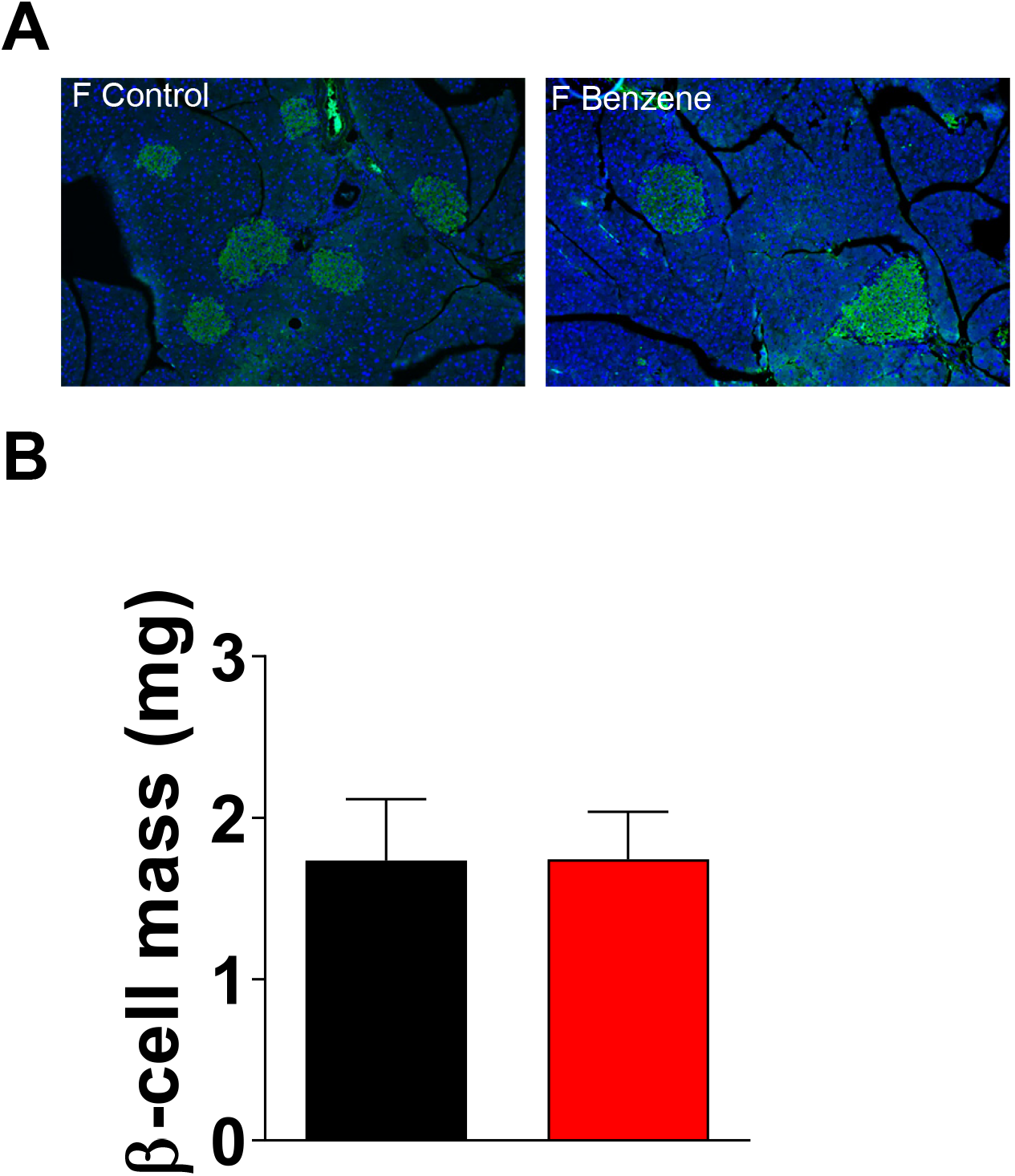

**Supplementary Figure 2.**
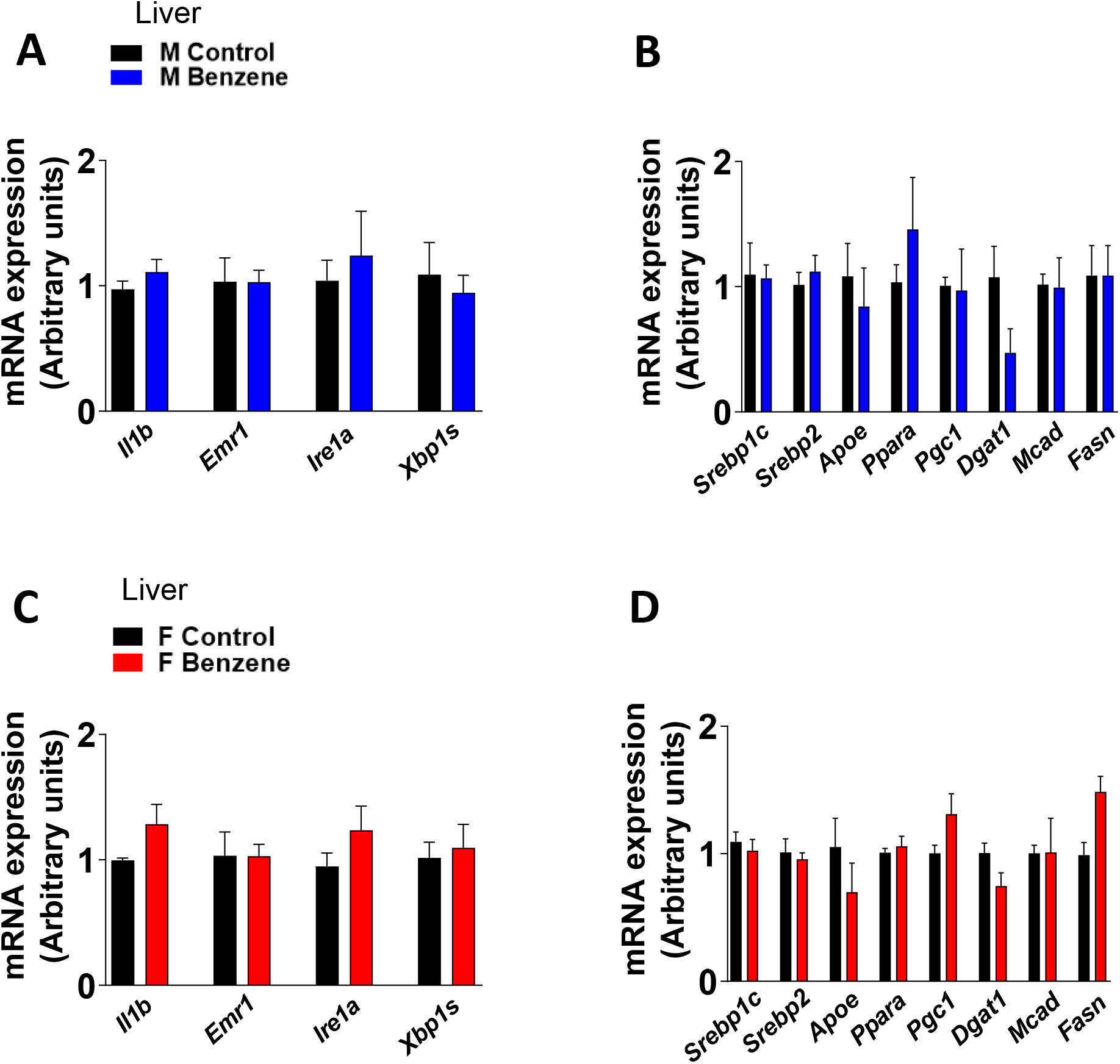

**Supplementary Table 1.**
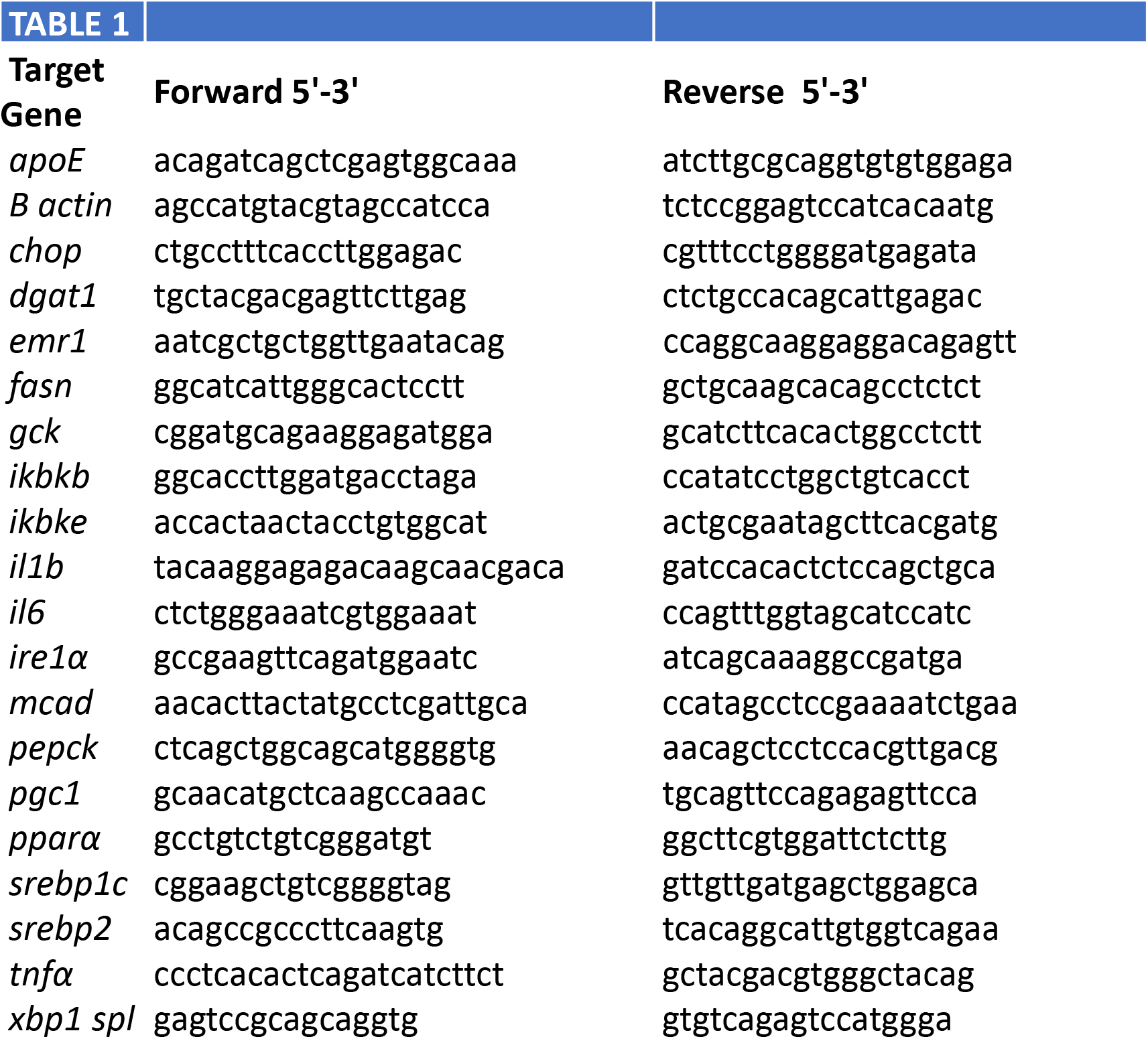

