## Supplemental for "Metabolic Reprogramming by *In Utero* Maternal Benzene Exposure"

### Supplementary Figure 1

**A**

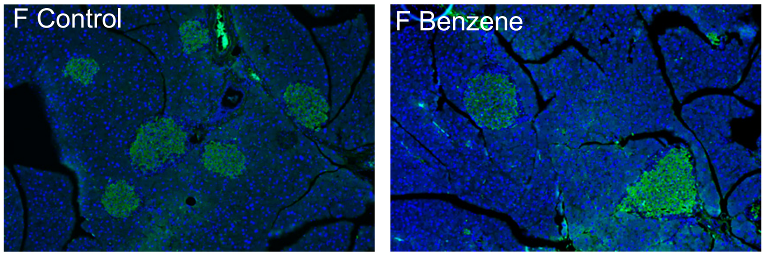


**B**

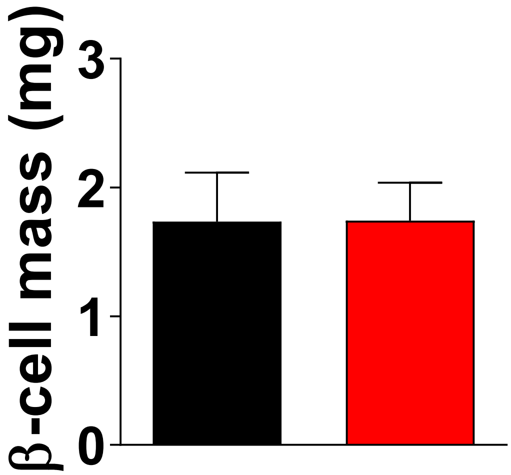

### Supplementary Figure 2

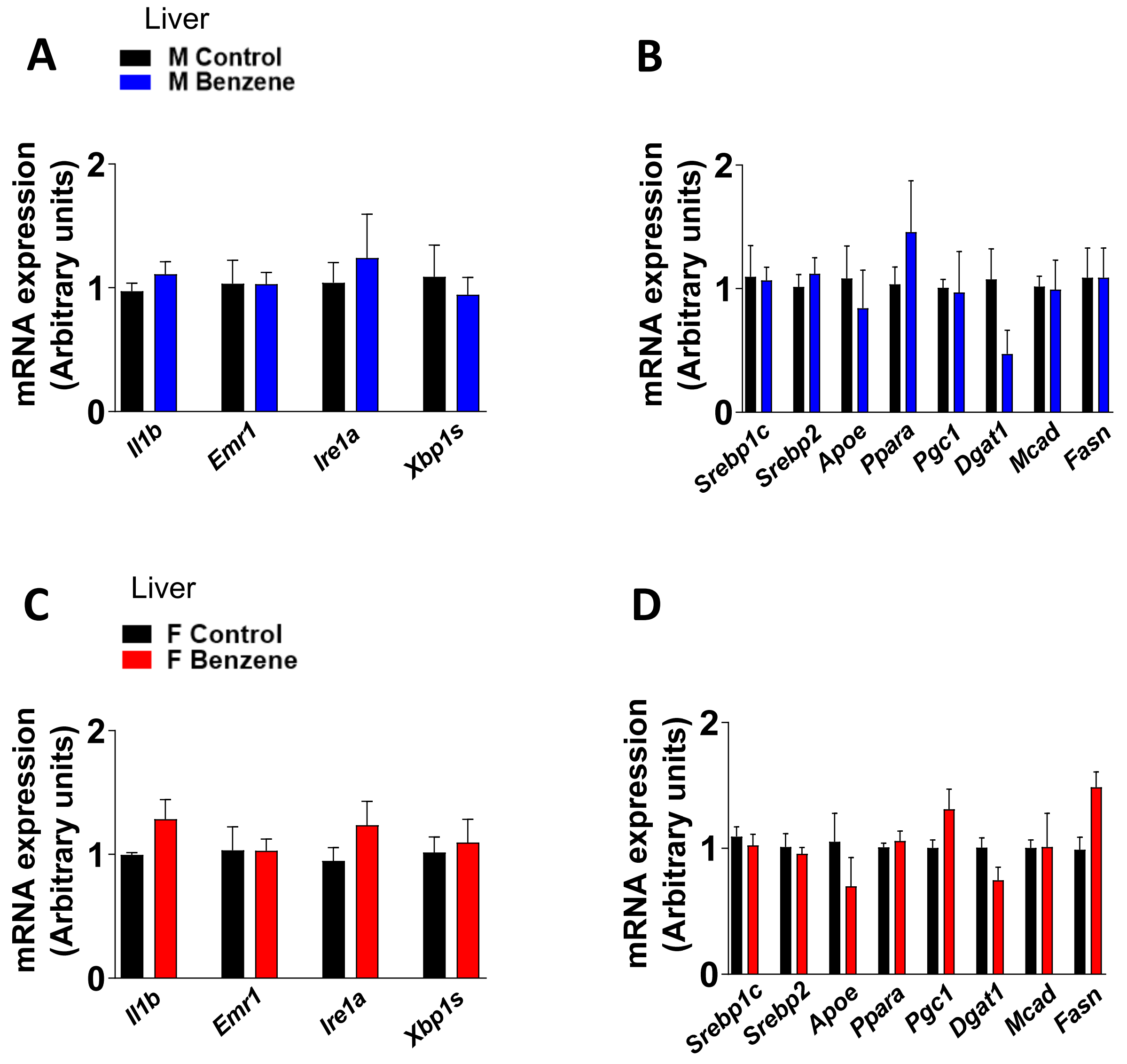

### Supplementary Table 1

| TABLE 1 | | |
|---|---|---|
| **Target Gene** | **Forward 5'-3'** | **Reverse 5'-3'** |
| *apoE* | acagatcagctcgagtggcaaa | atcttgcgcaggtgtgtggaga |
| *B actin* | agccatgtacgtagccatcca | tctccggagtccatcacaatg |
| *chop* | ctgcctttcaccttggagac | cgtttcctggggatgagata |
| *dgat1* | tgctacgacgagttcttgag | ctctgccacagcattgagac |
| *emr1* | aatcgctgctggttgaatacag | ccaggcaaggaggacagagtt |
| *fasn* | ggcatcattgggcactcctt | gctgcaagcacagcctctct |
| *gck* | cggatgcagaaggagatgga | gcatcttcacactggcctctt |
| *ikbkb* | ggcaccttggatgacctaga | ccatatcctggctgtcacct |
| *ikbke* | accactaactacctgtggcat | actgcgaatagcttcacgatg |
| *il1b* | tacaaggagagacaagcaacgaca | gatccacactctccagctgca |
| *il6* | ctctgggaaatcgtggaaat | ccagtttggtagcatccatc |
| *ire1α* | gccgaagttcagatggaatc | atcagcaaaggccgatga |
| *mcad* | aacacttactatgcctcgattgca | ccatagcctccgaaaatctgaa |
| *pepck* | ctcagctggcagcatggggtg | aacagctcctccacgttgacg |
| *pgc1* | gcaacatgctcaagccaaac | tgcagttccagagagttcca |
| *pparα* | gcctgtctgtcgggatgt | ggcttcgtggattctcttg |
| *srebp1c* | cggaagctgtcggggtag | gttgttgatgagctggagca |
| *srebp2* | acagccgcccttcaagtg | tcacaggcattgtggtcagaa |
| *tnfα* | ccctcacactcagatcatcttct | gctacgacgtgggctacag |
| *xbp1 spl* | gagtccgcagcaggtg | gtgtcagagtccatggga |
